# A novel nitrogen concentrating mechanism in the coral-algae symbiosome

**DOI:** 10.1101/2021.03.08.434475

**Authors:** Angus B. Thies, Alex R. Quijada-Rodriquez, Haonan Zhouyao, Dirk Weihrauch, Martin Tresguerres

**Affiliations:** Marine Biology research Division, Scripps Institution of Oceanography, University of California, San Diego, La Jolla, CA 92093, Unites States of America; Department of Biological Sciences, University of Manitoba, Winnipeg, MB, Canada

**Keywords:** Symbiosis, Rhesus channel, Ammonia, Coral Reef, Symbiodinium

## Abstract

Coral algal symbionts are hosted inside the symbiosome of gastrodermal cells, an intracellular compartment that isolates algae from the external environment and allows host cells to control the delivery of metabolites to their symbionts. However, the underlying molecular mechanisms are largely unknown. Here, we report the diel trafficking of NH_3_-transporting Rhesus (Rh) channels between the cytoplasm and the symbiosome membrane in the coral *Acropora yongei*, which matches established patterns of nitrogen delivery to endosymbionts. Heterologous expression in *Xenopus* oocytes established that *A. yongei* Rh (ayRhp1) is a channel that facilitates NH_3_ diffusion across membranes following its partial pressure gradient. Immunostaining revealed ayRhp1 is widely distributed throughout coral tissues and most abundantly present in oral ectodermal cells, desmocytes, and gastrodermal cells. In the latter, ayRhp1 was observed in the symbiosome membrane of alga-containing cells. Together with V-type H^+^-ATPases that make the symbiosome highly acidic (pH~4), ayRhp1 constitutes an NH_4_^+^-trapping mechanism analogous to that in mammalian renal tubule. Remarkably, ayRhp1 presence in the symbiosome membrane was higher during the day than the night. This indicates a regulatory mechanism that facilitates NH_4_^+^ delivery to alga during the day, likely to sustain high turnover rates of photosynthetic proteins, while restricting NH_4_^+^ delivery at night to maintain the endosymbiotic algae in a nitrogen-limited stage that stagnates their growth. The dynamic trafficking of proteins to and away from the symbiosome membrane is a previously unknown mechanism that contributes to metabolic regulation between symbiotic partners.

**Significance Statement:** The endosymbiotic relationship between corals and algae relies on the coordinated exchange of metabolites. Disruption of these metabolic exchanges can result in interruption of the symbiosis; however, the underlying molecular mechanisms are poorly understood. Here we report that *Acropora yongei* coral host cells express ammonia-transporting channel proteins (ayRhp1), which traffic to and away from the symbiosome membrane surrounding the endosymbiotic algae. In conjunction with the acidic symbiosome microenvironment, this mechanism allows host cells to regulate nitrogen delivery to endosymbionts sustaining essential functions while restricting growth. This work provides novel mechanistic information about metabolic regulation of animal-algae symbioses, and advances our understanding of physiological mechanisms that might determine coral local adaptation, resilience, and vulnerability to environmental stress including climate change.

## Introduction

Photosymbiotic associations between invertebrates and microalgae are widespread in aquatic environments. Perhaps the most well-known of these partnerships is that of reef-building corals (phylum: Cnidaria) and dinoflagellate symbiotic algae (family: Symbiodiniaceae), which is key to the evolutionary success of coral reef ecosystems (1). In an otherwise oligotrophic environment, the cnidarian host satisfies the majority of their energetic needs using photosynthates derived from their symbiont algae (2, 3). The host cells are believed to exercise considerable control over the metabolism of their symbionts, which favors both the production and release of algal photosynthates. This control is possible due to an architectural arrangement whereby coral gastrodermal cells host the algal symbionts intracellularly within an arrested phagosome known as the symbiosome (4–6). Because the symbiosome isolates the alga from the cytosol of the host cell, the symbiosome membrane necessarily mediates all metabolic exchanges between the symbiotic partners. In addition, the symbiosome membrane may serve as an interface for the coral to manipulate the alga’s microenvironment. For example, the coral symbiosome is markedly acidic (pH ~4) due to active H^+^ pumping by V-type H^+^-ATPases (VHAs) located in the symbiosome membrane (7). The acidic nature of the symbiosome drives CO_2_ accumulation as part of a carbon concentrating mechanism (CCM) that helps overcome the low affinity of algae Rubisco for CO_2_ thereby promoting algal photosynthesis (7). This H^+^ gradient has been proposed to additionally energize the movement of other essential nutrients and metabolites into or out of the symbiosome including nitrogen, phosphorous, and sugars (4, 7, 8). However, no additional molecular players or regulatory mechanisms have been definitely identified to date.

The vast majority of the symbiotic algae’s nitrogen demand is supplied by protein catabolism by their animal host, which produces waste as ammonia gas (NH_3_) and ammonium ion (NH_4_^+^) [collectively referred to as “total ammonia” (Tamm)] (3, 9). Indeed, rather than excreting its nitrogenous waste into the environment like most other aquatic animals (10), the coral symbiosis recycle Tamm via the glutamine synthase/glutamate dehydrogenase/glutamine oxoglutarate aminotransferase pathways (GS/GDH/GOGAT) (11–13). In addition, corals are able to take up NH_4_^+^ from seawater and transport it to their algal symbionts (14), and isolated algal symbionts take up and utilize NH_4_^+^ (15). Moreover, coral host cells are known to regulate Tamm delivery to their symbionts, and as a result the algae accumulate significantly more nitrogen in the light than in the dark (14, 16). The diel regulation of Tamm delivery by corals allows for host-control over the carbon and nitrogen metabolisms of symbionts (17) and, by extension, the growth rate and biomass of the symbiont population to prevent symbiont overgrowth that would disrupt the symbiosis (18). However, the mechanisms that mediate and regulate nitrogen transport to symbionts across the symbiosome membrane remain unknown.

Importantly, NH_3_ and NH_4_^+^ exist in pH-dependent equilibrium with pKa ~ 9.25, and accordingly >96% of Tamm is found as NH_4_^+^both in seawater (pH ~8) and in coral host cells (pH ~7.4) (19). However, the much lower pH in the symbiosome space has three critical and interlinked implications: first, a virtually nil NH_3_ partial pressure (*p*NH_3_) in the symbiosome space that should drive NH_3_ gas diffusion from the host cytoplasm; second, the immediate “trapping” of NH_3_ as NH_4_^+^ in the symbiosome space, which can be taken up by the alga thus maintaining the inwardly-directed NH_3_ diffusive gradient; and finally, an unfavorable electrochemical gradient for NH_4_^+^ transport into the symbiosome.

But despite being a gas, NH_3_ has limited permeability through lipidic membranes due to its strong dipole moment that makes it a polar molecule [reviewed in (20)]. In some plant-bacteria symbioses, NH_3_ transport across the symbiosome membrane is facilitated by nodulin-intrinsic proteins (21, 22); however, this protein family is exclusive to plants. Additionally, NH_3_ diffusion across biological membranes can be significantly enhanced by Rhesus (Rh) channels, a family of evolutionary conserved proteins present in eubacterial, invertebrate, and vertebrate lineages (23–27). Based on the observed upregulation of a Rh-like mRNA transcript upon establishment of symbiosis in anemones (28, 29), Rh channels have been suggested to play important roles in cnidarian-algae symbioses. However, the Rh-like mRNA was expressed in many soft coral cell subtypes (30), and therefore the coded protein probably plays multiple physiological roles. In addition, Rh channels are typically present in the cell outer plasma membrane [reviewed in (20, 31)], and few studies have localized Rh-like proteins to intracellular compartments or organelles (32, 33). Thus, the presence of coral Rh-like proteins in the symbiosome membrane cannot be taken for granted. Finally, some mammalian Rhs may transport both NH_3_ and NH_4_^+^ (20) and others have structural functions instead of facilitating Tamm transport across membranes (34–37). However, detailed studies about NH_3_ vs NH_4_^+^ transport capabilities by ‘primitive’ Rh proteins from invertebrate animals (termed “Rhp”) are very scarce. As a result, assessing the physiological role of the coral Rh-like coded protein and its potential involvement in delivering Tamm to algal endosymbionts requires elucidating their actual function as well as its cellular and subcellular localizations.

Given that NH_3_ diffusion through biological membranes is limited, we hypothesized that corals utilize Rh-like proteins to deliver NH_3_ to their algae across the symbiosome membrane, which would subsequently get trapped as NH_4_^+^ in the acidic symbiosome. To investigate this possibility, we cloned a Rh-like gene from the coral *Acropora yongei* (ayRhp1) and determined its phylogenetic relationship to other Tamm-transporting proteins. Then, we heterologously expressed ayRhp1 protein in *Xenopus* oocytes and measured Tamm transport under a range of pHs to determine whether it is a functional NH_3_ channel. Using custom made antibodies and immunocytochemistry, we observed that ayRhp1 protein was present in several cell types, including algae-containing gastrodermal cells. Furthermore, confocal super-resolution microscopy definitely determined the presence of Rhp1 in the symbiosome membrane. Notably, the proportion of cells with ayRhp1 in the symbiosome membrane doubled during the day compared to the night. These results suggest that trafficking of *A. yongei* Rh in and out of the symbiosome membrane during diel cycles contributes to the mechanism that controls the delivery of Tamm to algal symbionts in a diel manner.

## Results

### Rhp1 genes are widespread in corals

The cloned ayRhp1 cDNA open reading frame contains 1440 base pairs encoding a protein with a predicted molecular weight of 51.8 kDa. BLAST searches in genomic and transcriptomic databases revealed predicted ayRhp1 homologs in multiple coral species from both the robust and complex clades, which diverged from each other 300-400 million years ago (38). These coral Rh proteins clustered together with Rhp1 genes from invertebrate animals (Fig. S1).

The protein structure of ayRhp1 is similar to that of well-studied Rh channels from mammals (Fig. 1). It has 12 transmembrane helices and a N-linked glycosylation site (N61), which differentiate all animal Rh50 channels capable of Tamm transport (Rhag-cg, Rhp1-2) from the Rh30 proteins involved in structural functions (32). Crystallography and simulation studies have identified several key amino acid residues that are required for NH_3_ transport across mammalian RhCG (26, 27): a phenylalanine gate (F130, F235) and a cytosolic shunt (L193, T325, L328, I334, N341, N342) which recruit NH_4_^+^ at the external and internal vestibules, respectively, twin histidines that deprotonate NH_4_^+^ to NH_3_ (H185, H344), two highly-conserved aspartic acid residues that help shuttle the H^+^ back to the original compartment (D177, D336), and a hydrophobic transmembrane channel that selectively conducts NH_3_ but not NH_4_^+^ (37, 39). An alignment of ayRhp1 with RhCG reveals that the phenylalanine gate (F147, F251), the twin histidines (H202, H364) and analogous aspartate residues (D195, D356) are all conserved in ayRhp1, while the cytosolic shunt and hydrophobic channel-lining residues are highly-conserved (~83%, ~70% respectively). The overall high conservation of these key structures suggests that ayRhp1 facilitates NH_3_ transport; however, the lack of detailed functional studies on any Rhp1 protein precludes a definite assessment.

**Figure 1:**
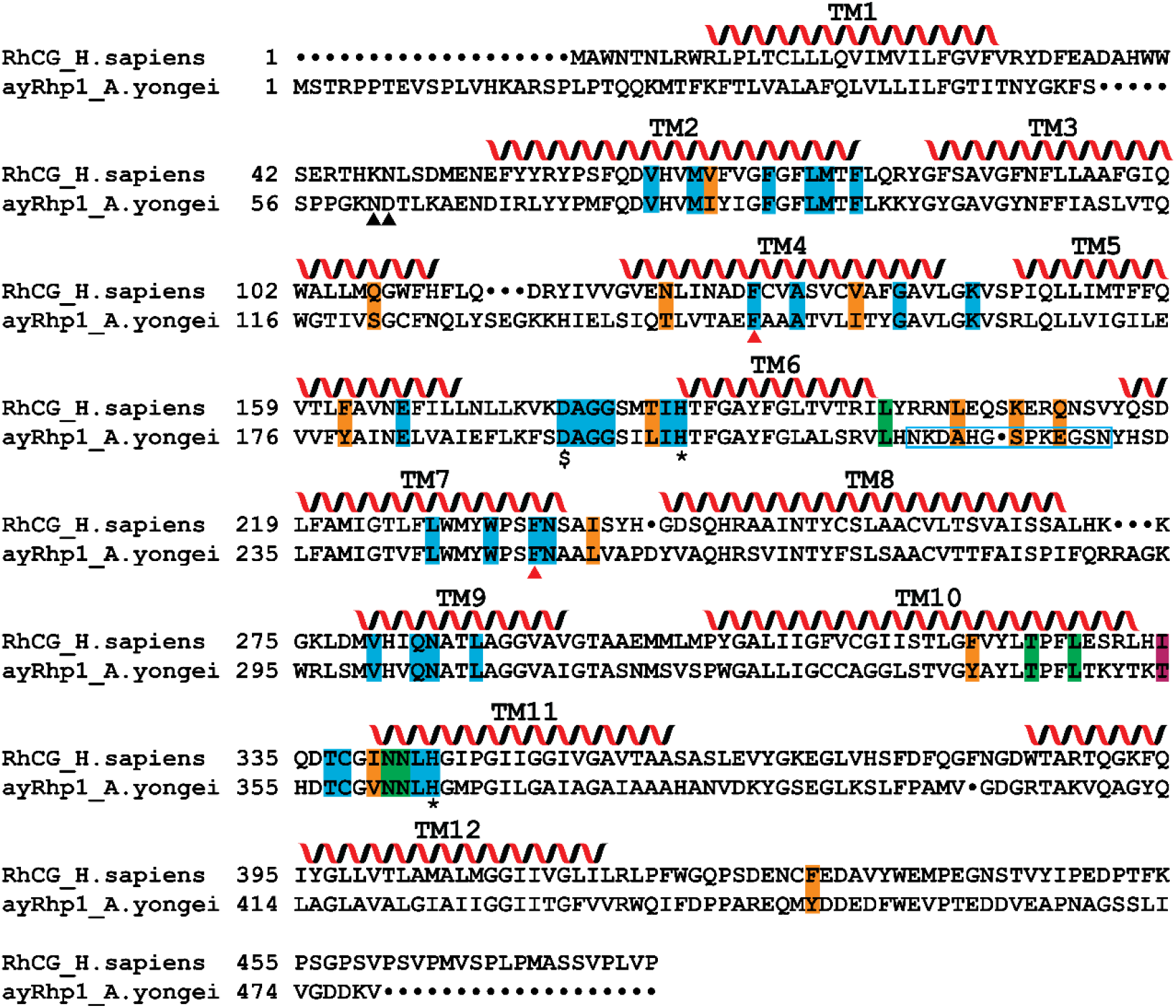
Alignment of ayRhp1 with *Homo sapiens* RhCG (NP_001307970.1). Conserved (blue) and mismatched (orange) hydrophobic channel-lining residues, conserved (green) and mismatched (magenta) cytosolic shunt residues, N-glycan site (black triangles), Phe-gate (red triangles), twin-Histidines (*), and the highly-conserved aspartate residues ($) as discussed in (26, 39) are marked. Gaps in sequences are denoted by black dots. The 12 transmembrane helices (TM), as predicted by TMHMM 2.0 (91, 92), for RhCG (26) and ayRhp1 are labeled with spirals.

### ayRhp1 Facilitates NH_3_ Transport

ayRhp1 was functionally characterized by measuring Tamm uptake rates in *Xenopus* oocytes injected with ayRhp1 cRNA. To avoid potential artefacts resulting from using radiolabeled [^14^C]-methylammonium as a Tamm analog (20), we used a hypochlorite-salicylate-nitroprusside based colorimetric assay to directly measure Tamm accumulation in oocytes and estimate Tamm uptake rate. The bath solutions contained 1 mM Tamm at pH 6.5, 7.5, or 8.5, resulting in 10-fold pKa-dependent [NH_3_] increases for every pH unit (1.8, 17.4, and 150.5 μM, respectively). Tamm uptake rate significantly increased from 10.2 ± 1.4 pmol Tamm min^−1^ at pH 6.5, to 36.9± 2.8 pmol Tamm min^−1^ at pH 7.5, and to 49.6 pmol Tamm min^−1^ at pH 8.5 (Fig. 2A), an apparent *Jmax* and *K_m_* of 51.9 and 7.1 pmol Tamm min^−1^, respectively (Fig. 2B). These results indicate that ayRhp1 transports NH_3_ following the partial pressure difference. Additionally, Tamm uptake rate of oocytes incubated in a solution with 10 mM Tamm at pH 7.5 was 50.3 ± 10.0 pmol Tamm min^−1^ (i.e. indistinguishable from the rate in the 1 mM Tamm pH 8.5 solution) (Fig. 2B). These two solutions have similar [NH_3_] (174.0 vs. 150.5 μM), but the former has >10-fold greater [NH_4_^+^] than the latter (9,826.0 μM vs. 849.5 μM). Altogether, these results established that ayRhp1 can facilitate NH_3_ diffusion following pH-dependent partial pressure gradients, and that Tamm transport is not directly dependent on the [NH_4_^+^] difference.

**Figure 2:**
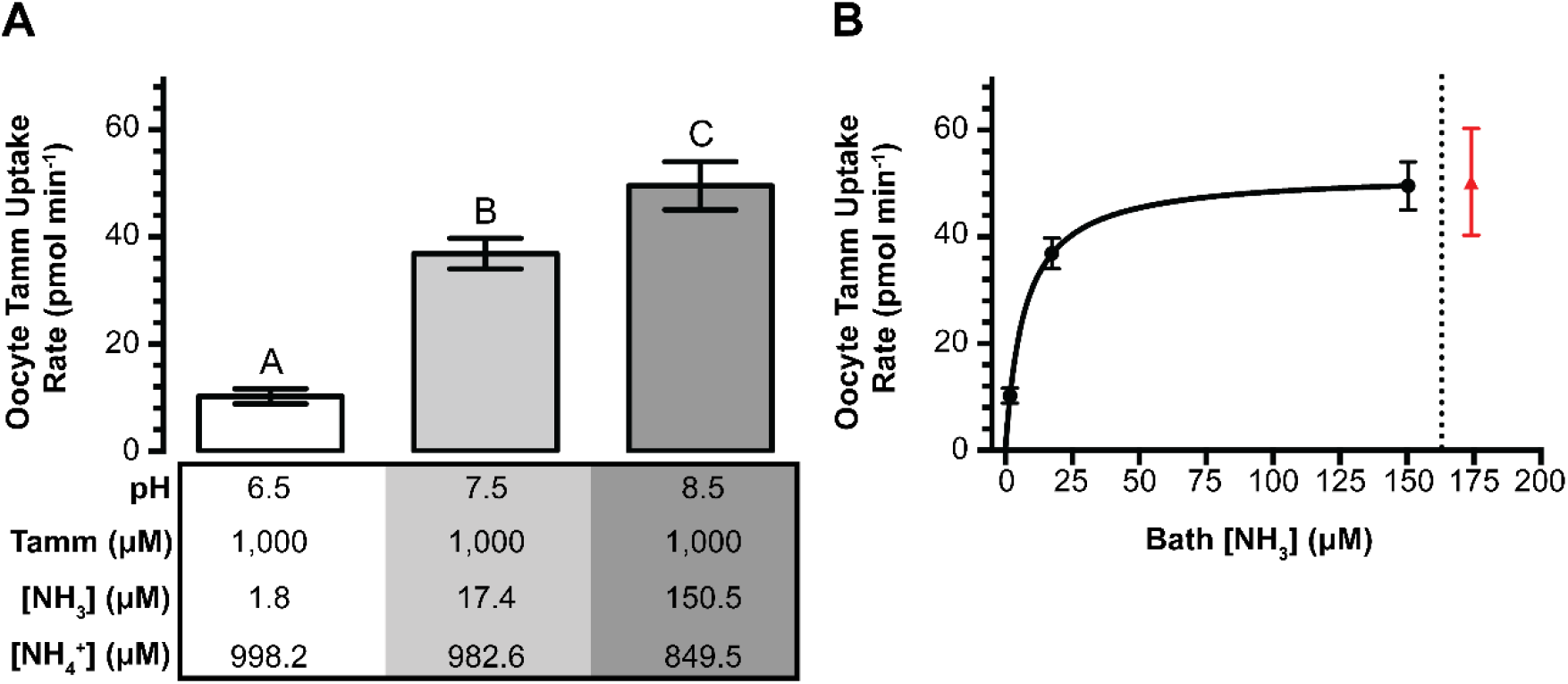
Functional characterization of ayRhp1. **(A)** Effect of [NH_3_] on total ammonium (Tamm) uptake rate in Xenopus oocytes expressing ayRhp1. Control Tamm uptake rates have been subtracted. Data shows mean ± S.E.M. of 6-8 oocytes; the letters denote significant differences (one-way ANOVA followed by Tukey’s multiple comparisons test; pH 6.5 vs. 7.5: p<0.0001, pH 7.5 vs. 8.5: p=0.0222, pH 6.5 vs. 8.5: p<0.0001). **(B)** Michaelis-Menten Tamm uptake kinetics calculated from the data shown in A (black dots to the left of the dotted line). Apparent Jmax and Km were 51.9 and 7.1 pmol Tamm min^−1^, respectively. The red triangle indicates Tamm uptake rate obtained in a solution with 10 mM Tamm at pH 7.5 (175 μM NH_3_ and 9.825 mM NH_4_^+^) (i.e. similar [NH_3_] to the previous data point, but ~10-fold higher [NH_4_^+^]).

### ayRhp1 protein is present in multiple coral cell types

Immunofluorescence microscopy using custom-made specific antibodies revealed high ayRhp1 protein expression throughout coral tissues (Fig. 3A). In the oral ectoderm, ayRhp1 was present in the apical membrane of columnar cells along the seawater-coral interface (Fig. 3B_1_). Although corals recycle most of their nitrogen waste through their algal symbionts, they also excrete some Tamm to the environment (3, 11). Thus, we hypothesize that ayRhp1 in ectodermal cells aids in nitrogenous waste excretion as previously described in gills and skin from fish and aquatic invertebrates (40, 41). Moreover, Tamm excretion may be facilitated by stirring of the boundary layer by ciliary beating, akin to mussels and polychaetes (40, 42). Notably, the surface of the oral ectoderm may host an extensive microbiome including nitrogen-fixing bacteria (43), and thus ayRhp1 may additionally facilitate the uptake of bacterially-fixed Tamm into the coral cells.

**Figure 3:**
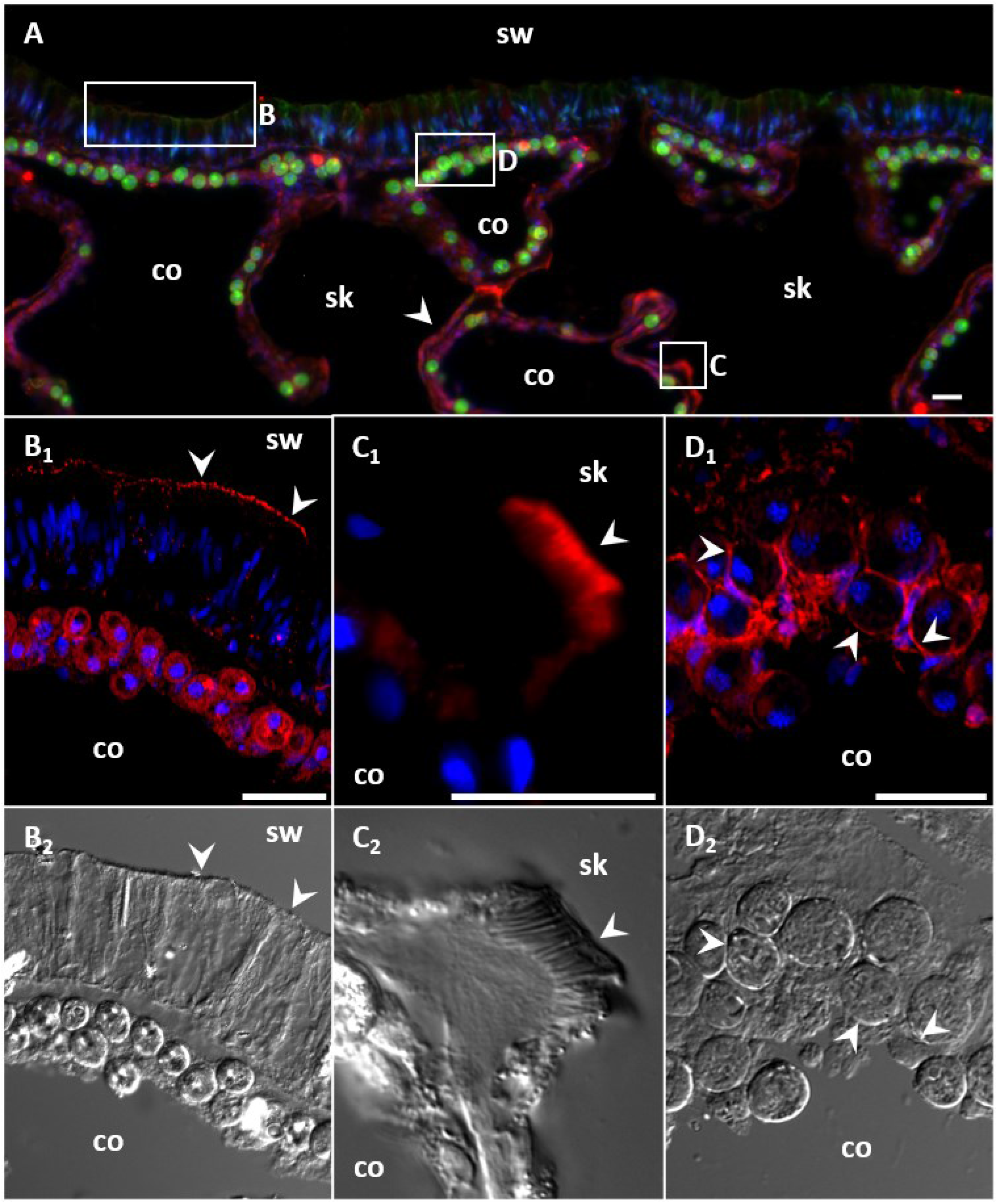
Immunolocalization of ayRhp1 in *A. yongei*.**(A)**Overview of *A. yongei* tissues; the boxes indicate regions of interest shown at higher magnification below, and the white arrowhead indicates ayRhp1-labeled calcifying cells. **(B_1_)**Apical membrane of columnar cells in the oral ectoderm. **(C_1_)**A desmocyte with intense signal in its apical region. **(D_1_)**Alga-containing gastrodermal cells. **(B_2_, C_2_, D_2_**) Corresponding brightfield differential interference contrast images; the white arrowheads mark corresponding locations in panels B, C, D. Nuclei are shown in blue, endogenous green fluorescent protein and chlorophyll are shown in green, and ayRhp1 immunofluorescence is shown in red. Red algal autofluorescence remains visible in panel B. sw: seawater, co: coelenteron, sk: skeleton. Scale bar = 20 μm.

In the calicodermis, ayRhp1 was expressed in both calcifying cells that deliver dissolved inorganic carbon, Ca^2+^, and matrix proteins for skeletal formation and in desmocytes that anchor living coral tissue to the skeleton. The ayRhp1 signal in desmocytes was very intense, especially at the apical membrane adjacent to the skeleton (Fig. 3C_1_, Fig. S3B_1_). Previous studies have provided morphological descriptions of coral desmocytes (44–47); however, to our knowledge this is the first description of any protein specifically expressed in this cell type. Interestingly, metabolic NH_3_ has been proposed to promote biological calcification of avian egg shells (48), land snail shells (49), and coral skeletons (50) by buffering H^+^ produced during CaCO_3_ precipitation. Whether ayRhp1 plays a role in the coral calcification mechanism is a topic for future studies. In the gastroderm, ayRhp1 was highly expressed in alga-hosting coral cells surrounding the symbiotic algae (Fig. 3D_1_), in a pattern that resembled that of VHA in the symbiosome membrane (7).

### ayRhp1 is present in the symbiosome membrane

The subcellular localization of ayRhp1 in alga-hosting coral cells was further studied by immunostaining isolated cells, as done previously for VHA (7). The ayRhp1 immunofluorescent signal was present around the algal cell (Fig. 4 A_1,2_-C_1,2_). Free algal cells released during the isolation procedure, identified by the lack of an adjacent host nucleus, did not have ayRhp1 signal (Fig. 4D_1,2_). In most cells, ayRhp1 was present in the thin region between the host cell nucleus and the alga (Fig. 4A_1,2_). Since these cells are tightly packed, this region is occupied by the symbiosome membrane (7, 51). The localization of ayRhp1 in the symbiosome membrane was further confirmed in cells containing two algae (Fig. 4B_1,2_), which contain a larger cytoplasmic region thus allowing for better discrimination of subcellular compartments. We also identified cells that had ayRh1 signal around the host cell nucleus rather than in between the nucleus and the alga (Fig. 4C_1,2_), which suggested ayRhp1’s presence in the cytoplasm or plasma membrane of the host cell.

**Figure 4:**
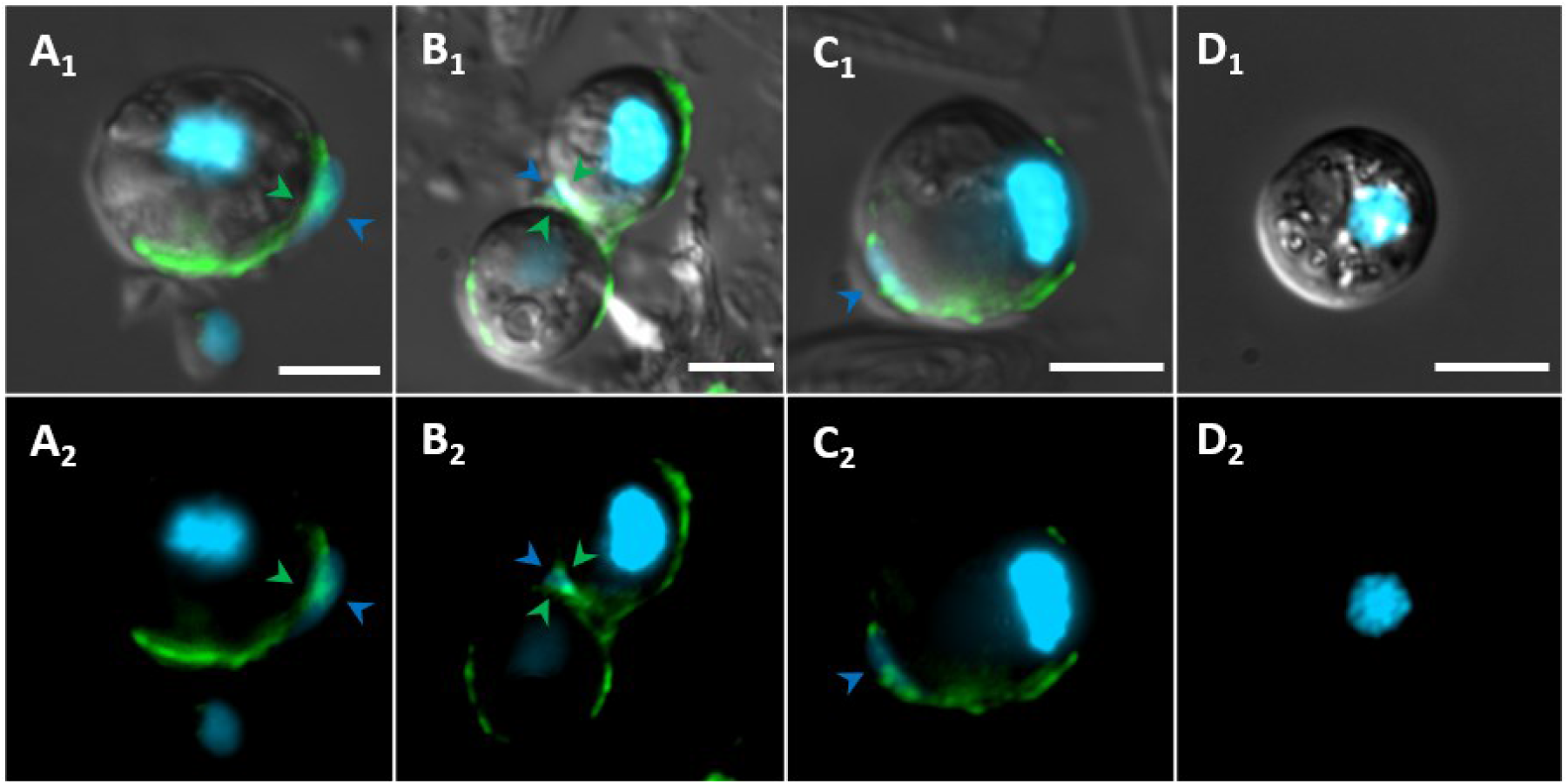
Epifluorescence immunolocalization of ayRhp1 in isolated alga-hosting gastrodermal cells. (**A_1_**) A coral host cell with ayRhp1 on the symbiosome membrane. (**B_1_**) A coral host cell containing two algal symbionts with ayRhp1 on both symbiosome membranes. (**C_1_**) A coral host cell with ayRhp1 on the exterior and sides of the host nucleus. (**D_1_**) Algal symbiont isolated from its host cell. (**A_2_, B_2_, C_2_, D_2_**) Corresponding images without brightfield differential interference contrast. Blue arrowheads mark nuclei of host cells; green arrowheads mark ayRhp1 symbiosome localization. Nuclei are shown in blue and ayRhp1 immunofluorescence in green. Scale bar = 5 μm.

Confocal super-resolution microscopy and co-immunostaining of ayRhp1 and Na^+^/K^+^-ATPase (NKA) allowed us to definitely establish ayRhp1’s subcellular localization within the tightly packaged host cells in coral tissues. Consistent with its universal presence in the plasma membrane (52), NKA outlined the perimeter of all alga containing host cells (Fig. 5). Although the ayRhp1 signal also was at the cell periphery, it was internal to that of NKA in most cells. This was most readily evident in the region around the nucleus, where NKA was external and Rhp1 was internal, indicating their respective presence in the plasma and symbiosome membrane (Fig. 5A_2,3_). However, some cells lacked ayRhp1 signal in the region between the host nucleus and the algae, and instead colocalized with NKA in the host plasma membrane (Fig. 5B1-3).

**Figure 5:**
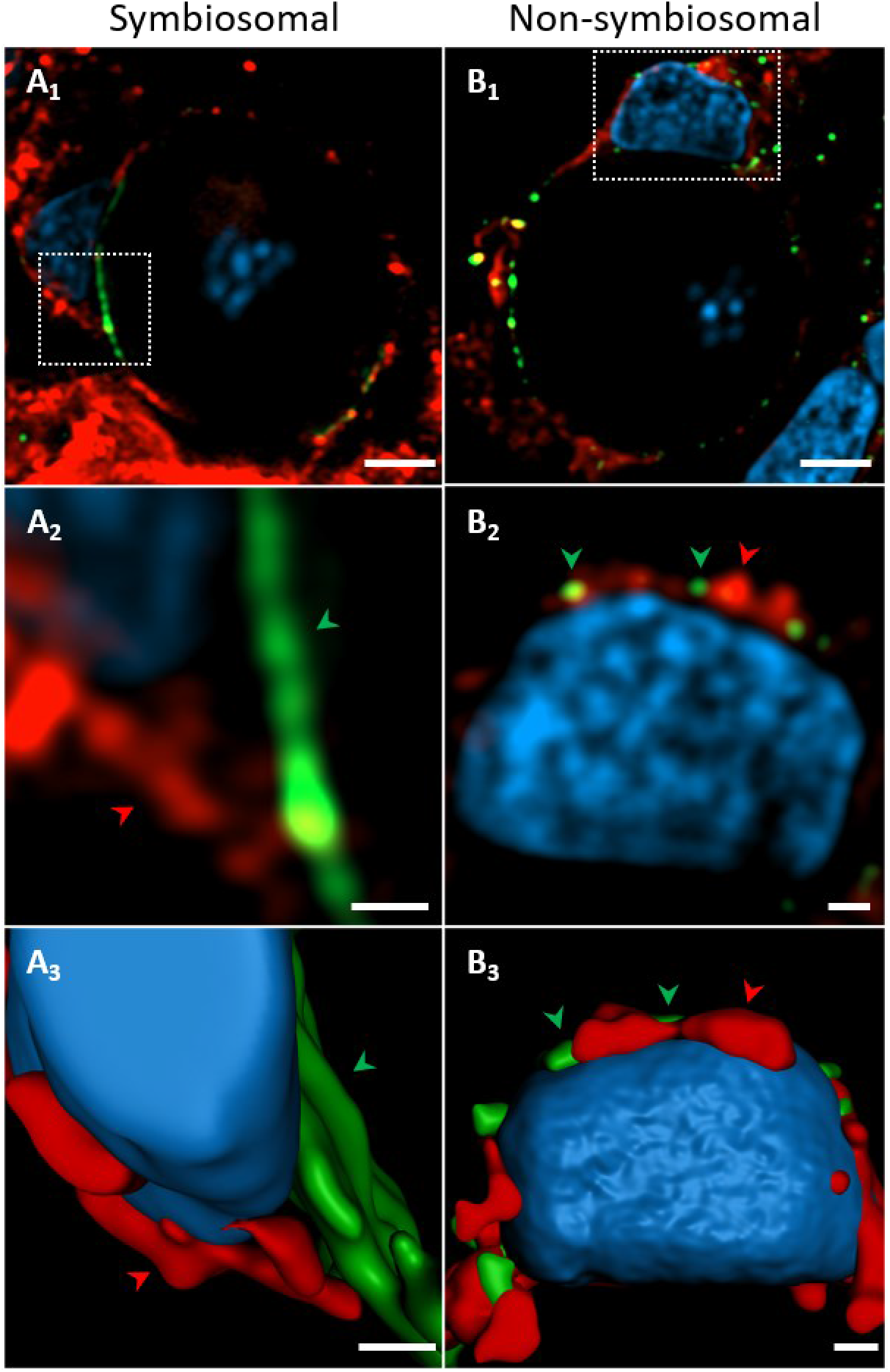
Super resolution confocal immunolocalization of ayRhp1 in alga containing coral cells (**A_1_, B_1_**) Cells displaying ayRhp1 in the symbiosome or plasma membrane, respectively (**A_2_, B_2_**) Higher magnification of the region denoted by the white boxes. (**A_3_, B_3_**) 3D renderings of A_2_, B_2_. Nuclei are shown in blue, ayRhp1 in green, and NKA in red. Notice the colocalization of ayRhp1 and NKA in B_2,3_. Scale bar = 5 μm, 0.5 μm, and 0.5 μm, respectively.

### Diel trafficking of ayRhp1 between the symbiosome and plasma membrane

In search for a reason behind the two distinct ayRhp1 subcellular localizations, we observed the subcellular localization of ayRhp1 over a diel cycle. To achieve sufficient replication (50 cells from 18 coral branches, three coral branches at each of six timepoints, for a total of 900 cells observed in blind fashion), we performed these observations on isolated alga-containing coral cells, as super resolution confocal microscopy was prohibitive time-wise. Based on the results shown in Fig. 4 and 5, cells with ayRhp1 signal in between the host cell nucleus and the alga were classified as “symbiosomal ayRhp1 localization”, and those lacking signal in that region were classified as “host plasma membrane localization”. Remarkably, the percentage of cells displaying ayRhp1 symbiosomal localization was significantly higher during the day, with a maximum of 61.3 ± 4.4% cells displaying this pattern at 13:00 h in contrast to only 26.0 ± 2.0% of cells at 23:00 h (*p* < 0.001). The percentage of cells displaying non-symbiosomal ayRhp1 signal followed the reciprocal pattern (Fig. 6). These results indicate that ayRhp1 is preferentially present in the symbiosome membrane during the day and in the host cell plasma membrane at night. To our knowledge, this is the first report of diel changes in proteomic makeup of the cnidarian symbiosome membrane and furthers the notion that this interface that separates symbiotic partners can be dynamically modified by the host cell to control the physiology of the alga.

**Figure 6:**
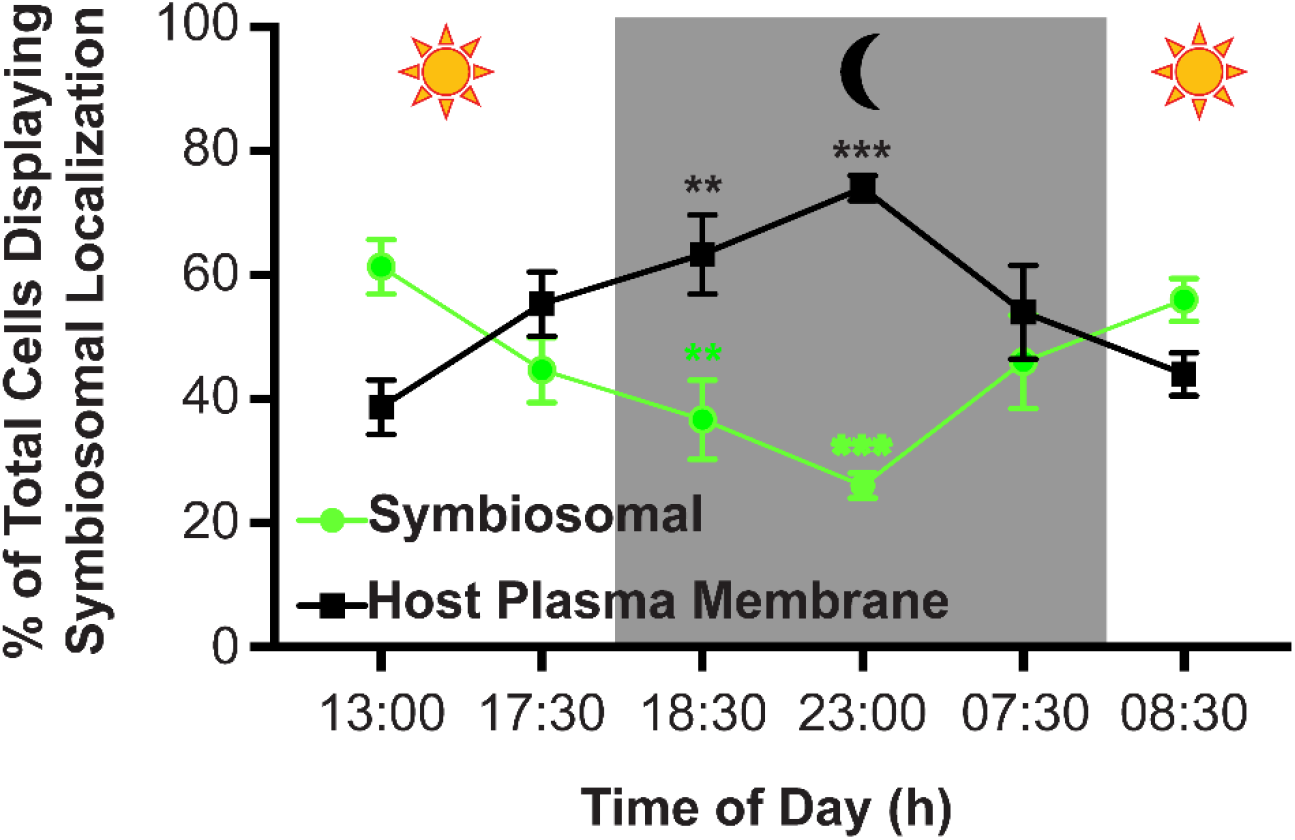
Percentage of total alga-containing host cells with symbiosomal ayRhp1 over a diel cycle. Data shows mean *±* S.E.M. n = 3 per timepoint, 50 cells per n, 900 cells total. The asterisks indicate significant differences with the 13:00 h timepoint (2-way repeated measures ANOVA followed by Dunnett post-test; *\*\*p<0.01; ***p<0.0001*).

## Discussion

The pKa for Tamm combined with the pH difference between the host cell’s cytosol and the symbiosome dictates >2,000 higher *p*NH_3_ in the former. Although this establishes a steep partial pressure gradient favoring NH_3_ diffusion into the symbiosome, diffusion across lipidic membranes is generally limited [reviewed in (20)]. The presence of ayRhp1 in the symbiosome membrane is poised to overcome this limitation thus enhancing NH_3_ delivery to the algal symbionts, in analogous manner to nodulin-intrinsic proteins in plant-*Rhizobium* symbioses (21, 22). Once inside the highly acidic symbiosome, NH_3_ will be immediately converted into NH_4_^+^, which cannot move across the plasma membrane, or through Rhp1 (Fig. 2B). This mechanism is known as “NH_4_^+^ acid-trapping, and is well-documented in diverse excretory epithelia from humans (26), teleost fishes (41), cephalopod and bivalve mollusks (40, 53), and crustaceans [reviewed in (31), (26, 40, 41, 53, 54)]. Moreover, NH_4_^+^ is the alga’s preferred nitrogen source (15), so algal NH_4_^+^ uptake will ensure the continuous conversion of NH_3_ into NH_4_^+^ in the symbiosome, which in turn will maintain NH_3_ diffusion from the host cytosol. By analogy to the CCM that facilitates symbiont photosynthesis (7), NH_3_ transport by ayRhp1 coupled to acid-trapping of NH_4_^+^ in the symbiosome can be considered a host-controlled nitrogen concentrating mechanism (NCM). The high degree of conservation amongst cnidarian Rh channels (Fig. 1) and the presence of an acidic symbiosome in anemones and corals from both the complex and robust clades (7) suggest that Rh-mediated NCMs is widespread in cnidarians. However, species- and environment-specific differences in the NCM contribution to Tamm transport are likely to exist and must be explored.

The increased proportion of cells displaying symbiosomal ayRhp1 localization during the day matches established patterns of increased Tamm delivery from host to symbiont (55) and symbiont nitrogen assimilation (14) during light conditions. The diurnal nitrogen supply is primarily used not to advance growth, but to sustain a high turnover of photosystem proteins and pigments damaged by UV radiation and electron transfer, which is essential for continued and efficient photosynthesis (14, 56, 57). This situation highlights the need for unique regulatory mechanisms in photosymbiotic associations compared to symbioses with non-photosymbiotic microbes, such as those in plant roots. Conversely, the removal of ayRhp1 from the symbiosome membrane at night would serve to restrict nitrogen supply to symbiotic algae, thus limiting the synthesis of non-photosynthetic proteins that would be essential to sustain their growth and reproduction (57, 58). This mechanism gains additional significance when we consider that the coral symbiosome is highly acidic in both light and dark conditions (7), and this implies a continued steep inwardly directed *p*NH_3_ gradient. Moreover, alga-containing gastrodermal cells are in contact with the gastrovascular cavity, or coelenteron. This compartment contains Tamm at concentrations that can be several hundred-fold higher compared to seawater (59), and experiences steep diel pH fluctuations that can reach up pH values as high as 9 during the day and as low as 6.75 at night (59–61). The preferential presence of ayRhp1 in the host plasma membrane at night would facilitate the removal of NH_3_ from the host cell into the coelenteron where it would be trapped as NH_4_, further restricting nitrogen supply to the algae at night.

The regulation of nitrogen delivery via changes in ayRhp1 subcellular localization is not mutually exclusive with regulation via the GS/GDH/GOGAT pathway (16, 62, 63), and in fact they complement each other. Indeed, the involvement of this pathway is largely based on changes in gene expression or enzyme activity upon transitioning from symbiotic to aposymbiotic stages (28) or during long term environmental disturbances (64, 65). But despite diel mRNA expression patterns (66), the abundance of most metabolic enzymes in coral cells, including that of GS, do not seem to change on a diel basis (67). However, this does not preclude diel regulation of enzyme activity by posttranslational modifications or substrate availability that could act synergistically with ayRph1 to control nitrogen to symbiotic algae. Moreover, ayRhp1 subcellular localization was not identical in all cells at any time period, indicating finer regulation based on position on the coral colony, symbiotic stage, or some other unidentified factors.

The mechanisms that regulate the translocation of ayRhp1 between the symbiosome and the plasma membrane are currently under investigation. The most likely candidate for mediating the redistribution of ayRhp1 is the cAMP pathway via the soluble adenylyl cyclase. This hypothesis is based on the established role of sAC in sensing changes in pHi in coral cells induced by algal photosynthetic activity (19) and matching diel changes in cAMP abundance in coral colonies (68). Furthermore, the sAC activity is known to trigger the redistribution of numerous ion-transporting proteins in a variety of cells types and organisms (8, 69–72). However, the putative involvement of sAC in regulating the redistribution of ayRhp1 in coral cells must be experimentally tested, which is not a trivial task. To put it in perspective, perhaps the best studied NH_4_^+^ excretion mechanism is in mammalian renal intercalated acid-secreting cells, which, like the coral symbiosome, relies on NH_4_^+^ acid-trapping mediated by Rh channels and VHA (73). Interestingly, renal Tamm transport can be modulated by changes in the subcellular distribution of Rh (74, 75). But despite the vastly larger research attention and resources devoted to biomedically-relevant topics compared to coral cell biology, the regulatory pathways underlying Rh redistribution in mammalian renal cells also remain unknown.

## Conclusions and perspectives

The ayRhp1 and VHA-dependent NCM identified here together with diel changes in ayRhp1 subcellular distribution provides a mechanism whereby coral host cells can supply nitrogen to their algal symbionts while still maintaining them in a nitrogen-limited state to control their growth. Interestingly, alterations in nitrogen delivery to coral symbiotic algae has been linked to several environmental stressors that result in disruption of the symbiosis at the colony level, commonly known as coral bleaching (76–81). Furthermore, the existence of more resilient species [such as *Porites ssp*. (82)] suggests species-specific differences in the mechanisms used to control Tamm delivery to symbionts, and identifying such differences may help predict differential vulnerability and resilience to bleaching. In addition, future studies must take into account that ayRhp1 is present in multiple cell types throughout coral tissues, which cannot be discerned using transcriptomics, proteomics, or metabolomics assays on bulk coral colony samples. With this in mind, techniques that allow the investigation of coral biology at the cellular and molecular levels such as nanoscale secondary ion mass spectrometry (‘nano-SIMS’) (16, 83) and confocal microscopy (84–87) are excellent (and essential) complements to “-omics” techniques. In particular, super resolution confocal microscopy will allow studying physiological processes at the coral-alga interface, the symbiosome membrane, in unprecedent detail.

## Materials and Methods

### Organisms

*A. yongei* colonies were maintained in flow-through seawater at 26°C with a 10/14h light/dark cycle with sunrise at 08:00 and sunset at 18:00. These coral colonies predominantly contain *Cladocopium sp* (formerly *Symbiodinium* clade C). See SI Methods for additional information on organism care.

### Cloning of ayRhp1

Following the methods of (19), total RNA was collected by flash-freezing a 2cm *A. yongei* nubbin in liquid nitrogen and crushing with a mortar and pestle into a fine powder. Powdered tissue was resuspended in TRIzol reagent (Invitrogen, Carlsbad, CA, USA) and total RNA was extracted following the manufacturer’s protocol. Total RNA was cleaned and concentrated using an RNeasy Plus Mini Kit (Qiagen, Hilden, Germany). cDNA was synthesized using SuperScript III Reverse Transcriptase (Invitrogen) and Oligo(dT) primers according to the manufacturer’s protocol. The resulting cDNA was used as template for all RT-PCR reactions. The full length ayRhp1 sequence was obtained following two rounds of RT-PCR using Phusion High Fidelity taq polymerase (New England Biolabs, Ipswitch, MA, USA) and NucleoSpin gel purification (Macherey-Nagel, Duren, Germany). The first round of RT-PCR used primers designed against untranslated regions of a predicted *Acropora digitifera* Rh mRNA (XP_015769291.1) (FWD primer 5’-CCACAATTCCGTC-3’, REV primer 5’-GTCCGAGACATCTTGCATACC-3’). In the second ‘nested’ round of RT-PCR, primers included oligonucleotide overhangs for In-Fusion Cloning (Clontech, Mountain View, CA, USA) into a pCR2.1-TOPO vector (Invitrogen) digested with EcoR 1 and EcoR V restriction enzymes. All cloned RT-PCR products were sequenced by Retrogen, Inc. (San Diego, CA, USA). The full ayRhp1 sequence can be found on Genbank (MH025799).

Protein sequences for the phylogenetic analysis were sourced from (88) and Genbank via BLASTtn search. Sequences were aligned using MUSCLE (89) and a maximum likelihood tree with 500 bootstraps was inferred by RAxML using a PROTGAMMA model of rate heterogeneity and a GTR substitution model (90). Accession numbers of all Rh sequences used in this analysis can be found in the supporting information file. Prediction of transmembrane helices was performed using TMHMM 2.0 (http://www.cbs.dtu.dk/services/TMHMM-2.0/) as per (91, 92).

### Antibodies

Custom-made, affinity purified anti-ayRhp1 rabbit polyclonal antibodies were developed (GenScript USA, Inc., Piscataway, NJ, USA) against the peptide HNKDAHGSHKEGSN, which is present in a putative Rhp1 protein predicted from the *A. digitifera* genome (XP_015769291.1) (93). This epitope has just one amino acid difference in ayRhp1 (HNKDAHGS**P**KEGSN). NKA was immunolocalized with a commercially available monoclonal antibody (SC-48345, Santa Cruz Biotechnology, Dallas, TX, USA).

### ayRhp1 Protein Expression and Antibody Validation

Using methods adapted from (7), *A. yongei* tissue was removed from the skeleton using an airbrush loaded with homogenization buffer. Briefly, homogenate was sonicated on ice and centrifuged to pellet down debris; the supernatant was kept on ice. Sample protein concentrations were determined using a Bradford Protein Assay (Bio-Rad, Hercules, CA, USA) with a bovine serum albumin standard curve. Samples were then incubated in 4x Laemmli sample buffer (Bio-Rad) and 10%β-mercaptoethanol before heating at 90°C for 3 min and loaded onto an SDS/PAGE gel. Following electrophoresis, proteins were transferred from the gel onto a PVDF membrane using a Mini Tans-Blot Cell (Bio-Rad) overnight. The membrane was blocked with 5% powdered fat-free milk in TBS-T for 1 h on a shaker at room temperature before overnight incubation on a shaker (4°C) with anti-ayRhp1 primary antibody (0.216 μg/ml), primary antibody with 400x excess peptide on a molar base (‘pre-absorption control’), or pre-immune serum (0.216 μg/ml) diluted in blocking buffer. Membranes were washed with 4 x 15 min TBS-T washes prior to incubation with secondary antibody (goat anti-rabbit-HRP diluted 1:10,000, Bio-Rad) for 1 h on a shaker at room temperature. Membranes were again washed with 4 x 15 min TBS-T washes and a final 15 min TBS wash prior to band development with an ECL Prime Western Blot Detection Kit (GE Healthcare, Chicago, IL, USA) and imaged using a Chemidoc Imaging system (Bio-Rad) (Fig. S2). See SI Methods for additional information.

### Plasmid Preparation and cDNA synthesis for *Xenopus laevis* Expression of ayRhp1

The open reading frame of ayRhp1 was amplified from pCR2.1 TOPO-ayRhp1 vector (see cloning of ayRhp1) using Q5 high fidelity DNA polymerase (New England Biolabs, Ipswich, MA, USA) and the restriction site-containing primers (forward primer, 5’-ATACCCGGGATGTCTACTCGACCTCCTACG -3’; reverse primer: 5’-GGCAAGCTTTTACACTTTATCATCTCCGAC -3’), subcloned by sticky-end ligation with T4 ligase (New England Biolabs) into XmaI and HindIII restriction sites of a PGEM-HE vector containing *Xenopus* beta globin 5’- and 3’-UTR sequences flanking the cloning site. Proper insertion was verified by multi-cutting restriction enzymes EcoR1, BamHI, and SphI, heat inactivation at 65 °C and column purification (GeneJet PCR Purification Kit; ThermoFisher Scientific, Waltham, MA, USA). The *in vitro* transcription of ayRhp1 capped mRNA (cRNA) was performed with HiScribe T7 ARCA mRNA kit (New England Biolabs) on SphI linearized pGEM-HE-ayRhp1 vector followed by column purification (RNeasy MiniElute Cleanup Kit; Qiagen). The cRNA was quantified spectrophotometrically (Nanodrop, ND-1000; Thermofisher Scientific) and its integrity was assessed on a denaturing MOPS agarose gel.

### Oocyte Microinjection

Stage VI-V oocytes were collected from mature female *Xenopus laevis* (94). Briefly, the frogs were euthanized via decapitation, and the ovary was dissected and placed in Ca^2+^-free oocyte ringer (OR2) solution (in mM: 82.5 NaCl, 2.5 KCl, 1 MgCl_2_, 1 Na_2_HPO_4_, 5 HEPES, pH 7.5) containing 1 mg ml^−1^ collagenase type VI (ThermoFisher Scientific). After incubation under gentle agitation for 90 min at room temperature, collagenase activity was terminated by rinsing the oocytes three times in OR2 containing 1 mM CaCl_2_. Oocytes were then manually sorted, rinsed, and allowed to recover in OR2 sterilized using vacuum bottle-top filters (EMD MilliporeTM SteritopTM), overnight at 16 °C (Fisherbrand Mini Refrigerated Incubator). Oocytes were injected with 18.4 ng of ayRhp1 cRNA (36.8 nL with 0.5 ng nl^−1^) (ayRhp1) or equivalent volume of nuclease-free water (control) using a Nanoject II or III auto-nanoliter injector (Drummond Scientific, Broomall, PA, USA). Experiments were conducted three days post-injection; during this time the oocytes were stored in OR2 supplemented with 2.5 mM sodium pyruvate, 1 mg ml^−1^ penicillin-streptomycin and 50 μg ml^−1^ gentamicin. Oocytes that died during experiments were discounted from analyses. All procedures followed the Guidelines of the Canadian Council on Animal Care and were approved by the University of Manitoba Animal Research Ethics Board.

### Oocyte Tamm Uptake Rates

Groups of 24 oocytes were placed in 15 mL tubes and incubated for 1 h at room temperature in OR2 solutions with: (a) 0 mM NH_4_Cl, pH 7.5; (b) 1 mM NH_4_Cl, pH 6.5; (c) 1 mM NH_4_Cl, pH 7.5, (d) 1 mM NH_4_Cl, pH 8.5; or (e) 10 mM NH_4_Cl, pH 7.5. Osmolarity was maintained by substituting NaCl with NH_4_Cl and pH was adjusted by adding NaOH or HCl. Following incubation, oocytes were washed in ice cold OR2 to remove excess NH_4_Cl and placed in groups of three oocytes in 27 μL 6% perchloric acid to deproteinize samples (95). After pH neutralization with 3 M KOH, samples were diluted 1:10-1:40 with MilliQ water and Tamm was measured using a hypochlorite-salicylate-nitroprusside based assay (96). Tamm uptake rate was calculated according to the formula

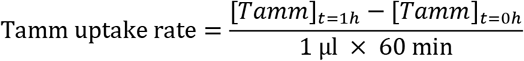
where [Tamm]t=1h is the Tamm measured in oocytes after incubation in OR2 b-e for 1h, [Tamm]t=0h is Tamm measured in oocytes in OR2 a prior to the start of the incubations, 1 μl is the average oocyte volume, and 60 min was used to calculate rates on a per minute basis. The average Tamm uptake rate of control oocytes (water injected) were subtracted from that of ayRhp1 oocytes prior to statistical analysis (Fig. S4). Tamm uptake kinetics were calculated using a non-linear regression to fit the Michaelis-Menten equation.

### Immunolabeling of ayRhp1 in Tissue Sections and Isolated Cells

Following (97), *A. yongei* nubbins and were fixed and decalcified for immunohistochemistry. Using methods adapted from (51, 97), isolated *A. yongei* cells were prepared for immunohistochemistry by submerging a nubbin in 0.2 μm filtered seawater and brushed with a toothbrush. Cells were collected via centrifugation and fixed. Fixative was removed via centrifugation and cells were resuspended in S22 buffer before plating on glass slides and immunolabeling.

Tissue sections and isolated cells were blocked for 1 h at room temperature in blocking buffer (4 ml PBS-TX, 80 μl normal goat serum, 0.8 μl keyhole limpet hemocyanin solution), followed by overnight incubation (4°C) with anti-ayRhp1 antibodies (2.16 μg/ml), anti-ayRhp1 antibodies pre-absorbed with excess peptide (8.64 μg/ml) or pre-immune serum (2.16 μg/ml) (all in blocking buffer) (Fig. S3). Slides were washed in PBS-TX to remove unbound anti-ayRhp1 antibodies (3 x 5 min). Secondary antibodies (goat anti-rabbit-Alexa Fluor555, 4 μg/ml in blocking buffer; Invitrogen) were then added for 1 h at room temperature followed by DAPI DNA Stain (1 μg/ml; Invitrogen) for 5 min at room temperature. Slides were again washed PBS-TX to remove unbound secondary antibody (3 x 5 min) and samples were imaged using a fluorescence microscope (Zeiss AxioObserver, Carl Zeiss AG, Oberkochen, Germany).

### Assessment of ayRhp1 Subcellular Localization over Day-Night Cycles

Cell isolations were prepared from coral nubbins randomly selected from three separate tanks. Nubbins were sampled 30 min before and after sunrise and sunset (07:30, 08:30, 17:30, 18:30 h) as well as halfway between lighting condition changes (13:00, 23:00 h). Samples taken during the day were continually illuminated during cell isolation and fixation while those taken during the night were kept in the dark. At each timepoint, cells were immonostained for ayRhp1 and imaged as described above. Starting from the upper-right corner of the field of view, the first 50 intact alga-hosting *A. yongei* cells displaying ayRhp1 signal were counted and classified into one of three subcellular localization patterns: “predominantly symbiosome localization” (the ayRhp1 signal formed a distinct and continuous ring in between the nuclei from the coral host cell and the algae), “predominantly cytosolic localization” (the ayRhp1 signal was diffuse throughout the coral host cell cytoplasm), and “mixed” (the ayRhp1 signal displayed a similar proportion of symbiosome and cytosolic localization). Cells were counted and classified in a double-blind manner: slides were named with random identifiers by an independent person before being observed on the microscope by another person. Classification was conducted during observation through the microscope eyepiece, as this allowed a better determination of ayRhp1 subcellular localization by rapid and repetitive adjustments to the fine focus and alternation between the Alexa Fluor555 and DAPI channels. Cells from three separate branches were classified at each timepoint, resulting in a total of 150 cells per timepoint and 900 cells in total. Timepoints were matched with random slide names only once all 900 cells were classified. To capture representative images of the two subcellular localization patterns, tissues labeled with ayRhp1 and NKA primary antibodies were observed on a super resolution confocal fluorescence microscope (Zeiss LSM 900 with Airyscan, Carl Zeiss AG) using goat anti-rabbit-Alexa Fluor488 and goat anti-mouse-Alexa Fluor568 secondary antibodies (4 μg/ml in blocking buffer; Invitrogen) and DAPI (1 μg/ml; Invitrogen). 3D reconstructions of z-stacks were generated using Imaris 9.0 (Bitplane, Zurich, Switzerland).

### Statistical Analysis

All statistical tests were run in GraphPad Prism 7 (San Diego, CA, USA). Ammonia flux and ayRhp1 localization data were tested for normality and homogeneity of variance using D’Agostino & Pearson or Shapiro-Wilk normality tests and Brown-Forsythe tests. Tamm uptake data were analyzed using one-way ANOVA with Tukey’s multiple comparisons. Data from ayRhp1 localization in isolated cells was analyzed using 2-way repeated measures ANOVA followed by Dunnett post-test using the data from 13:00h as control. Alpha was set at 0.05 for all statistical tests.

## Author Contributions

A.B.T, A.R.Q.R, D.W. and M.T. designed research; A.B.T., A.R.Q.R, and H.Z. performed research; A.B.T., A.R.Q.R, H.Z., D.W., and M.T. analyzed data; A.B.T. and M.T. wrote the paper.

## Acknowledgments and Funding Sources

We thank Michael Romero (Mayo Clinic) for his kind gift of *Xenopus* expression vector, Mikayla Ortega, Samantha Nöel, Amalia Serna, Cameron Hassabi, and Phil Zerofski (Scripps Institution of Oceanography) for their help in maintaining coral cultures, and Mikayla Nash (University of Manitoba) for support with oocyte sorting and maintenance. This work was partially supported by National Science Foundation (NSF) EF #1220641 to MT, NSF Graduate Research (GRFP 2019271478) and SIO Doctoral Scholar Fellowships to ABT, Natural Sciences and Engineering Research Council of Canada (NSERC) Postgraduate Doctoral Scholarship-Doctoral to ARQR, University of Manitoba Graduate Fellowship to HZ, and NSERC Discovery grant (RGPIN/5013-2018) to DW.

