## Supplemental Information for "A novel nitrogen concentrating mechanism in the coral-algae symbiosome"

**This PDF file includes:**

Supplementary Methods
Figures S1 to S4
Table S1
SI References

### Supplementary Methods Text
#### Organisms

*A. yongei* nubbins were originally sourced from the Birch Aquarium at Scripps and reared in a heated flow-through seawater aquaria at Scripps Institution of Oceanography for at least 1 year prior to the experiments. These coral colonies predominantly contain *Cladocopium sp* (formerly *Symbiodinium* clade C). Average [$PO_4^{3-}$], [$NO_3^-$], [$NO_2^-$] and [$NH_4^+$] were 0.35 ± 0.02 μM, 0.29 ± 0.11 μM, 0 μM and 0.54 ± 0.1 μM, respectively (Southern California Coastal Ocean Observing System; https://sccoos.org/harmful-algal-bloom/). Temperature was kept at 26°C, light/dark cycle was 10/14h with sunrise at 08:00 and sunset at 18:00 (LED Fixture lights, Orbit Marine, model 4103-B). Light intensity in the aquaria was measured to be 120 μE (MSC15 Spectral Light Meter, Gigahertz-Optik, Amesbury, MA, USA).

#### ayRhp1 Protein Expression and Antibody Validation

Using methods adapted from (1), *A. yongei* tissue was removed from the skeleton using an airbrush loaded with homogenization buffer. Homogenate was sonicated on ice for 4 x 10-sec bursts with 1 min rest in-between. The sonicated homogenate was then centrifuged (500 x g, 15 min, 4°C) to pellet down debris; the supernatant was kept on ice but not frozen. Sample protein concentrations were determined using a Bradford Protein Assay (Bio-Rad, Hercules, CA, USA). 4x Laemmli buffer (Bio-Rad) and 10% β-mercaptoethanol were added to samples before heating at 90°C for 3 min. 22.5 μg protein and 4 μl of Precision Plus Dual Color Protein Standards (Bio-Rad) were loaded into a 10% polyacrylamide SDS-PAGE gel in a Mini-Trans Blot Cell (Bio-Rad) with running buffer (25 mM Tris, 190 mM glycine, 0.1% SDS). Electrophoresis was run for 100 min at 100 V (4°C).

Following electrophoresis, the gel was washed in distilled water for 5 min and equilibrated in Towbin buffer (25 mM Tris pH 8.3, 192 mM glycine, 20% (v/v) methanol) for 15 min. Proteins were transferred from the gel onto a PVDF membrane using a Mini Tans-Blot Cell (Bio-Rad) overnight in Towbin buffer (0.09 A, 4°C). PVDF membranes were washed in Tris-buffered Saline + 0.1% Tween detergent (TBS-T) for 15 min on a shaker at room temperature to remove excess transfer buffer prior to blocking. Membranes were then blocked with 5% powdered fat-free milk in TBS-T for 1 h on a shaker at room temperature.

Following blocking, membranes were incubated overnight on a shaker (4°C) with anti-ayRhp1 primary antibody (0.216 μg/ml), primary antibody with 400x excess peptide on a molar base ('pre-absorption control'), or pre-immune serum (0.216 μg/ml) diluted in blocking buffer. Membranes were then washed with 4 x 15-min TBS-T washes prior to incubation with secondary antibody (goat anti-rabbit-HRP diluted 1:10,000, Bio-Rad) for 1 h on a shaker at room temperature. Membranes were again washed with 4 x 15-min TBS-T washes and a final 15-min TBS wash prior to band development with an ECL Prime Western Blot Detection Kit (GE Healthcare, Chicago, IL, USA) and imaged using a Chemidoc Imaging system (Bio-Rad).

**Immunolabeling of ayRhp1 in Tissue Sections and Isolated Cells**

Following (2), *A. yongei* nubbins were fixed for immunohistochemistry by immersion in S22 buffer (450 mM NaCl, 10 mM KCl, 58 mM $MgCl_2$, 10 mM $CaCl_2$, 100 mM Hepes, pH 7.80) supplemented with 4% paraformaldehyde overnight on a rocking platform at 4°C. Nubbins were then transferred to calcium-free S22 buffer supplemented with 0.5 M EDTA to decalcify the skeleton and with 0.5% paraformaldehyde to preserve tissue fixation; this decalcification buffer was changed daily for 2 weeks. Once decalcified, nubbins were dehydrated (50% ethanol for 5 h, 70% ethanol overnight, 95% ethanol for 20 min, 100% ethanol 3 x 20 min, SafeClear for 3 x 20 min), embedded in paraffin wax (3 x 30 min) and allowed to solidify for 48 h before microtome sectioning onto glass slides. *A. yongei* tissue sections were deparaffinized and serially rehydrated in SafeClear for 3 x 10 min, 100% ethanol for 10 min, 95% ethanol for 10 min, 70% ethanol for 10 min, and PBS with 0.2% (v/v) Triton-X-100 (PBS-TX) for 10 min.

Using methods adapted from (2, 3), isolated *A. yongei* cells were prepared by submerging a nubbin in 0.2 μm filtered seawater and brushed with a toothbrush for 1 min. Cells were collected via centrifugation (3,000 x g, 4 minutes, 4°C) and fixed by resuspension in S22 buffer with 4% paraformaldehyde for 15 min on ice. Fixative was removed via centrifugation (3,000 x g, 4 minutes, 4°C) and cells were resuspended in ~500μL of S22 buffer. Cells were then pipetted onto glass microscope slide and allowed to air dry at 4°C for no longer than 1 h before proceeding to immunolabeling.

Tissue sections and isolated cells were blocked for 1 h at room temperature in blocking buffer (4 ml PBS-TX, 80 μl normal goat serum, 0.8 μl keyhole limpet hemocyanin solution), followed by overnight incubation (4°C) with anti-ayRhp1 antibodies (2.16 μg/ml), anti-ayRhp1 antibodies pre-absorbed with excess peptide (8.64 μg/ml) or pre-immune serum (2.16 μg/ml) (all in blocking buffer) (Fig. S3). Slides were washed in PBS-TX to remove unbound anti-ayRhp1 antibodies (3 x 5 min). Secondary antibodies (goat anti-rabbit-Alexa Fluor555, 4 μg/ml in blocking buffer; Invitrogen) were then added for 1 h at room temperature followed by DAPI DNA Stain (1 μg/ml; Invitrogen) for 5 min at room temperature. Slides were again washed PBS-TX to remove unbound secondary antibody (3 x 5 min) and samples were imaged using a fluorescence microscope (Zeiss AxioObserver, Carl Zeiss AG, Oberkochen, Germany).

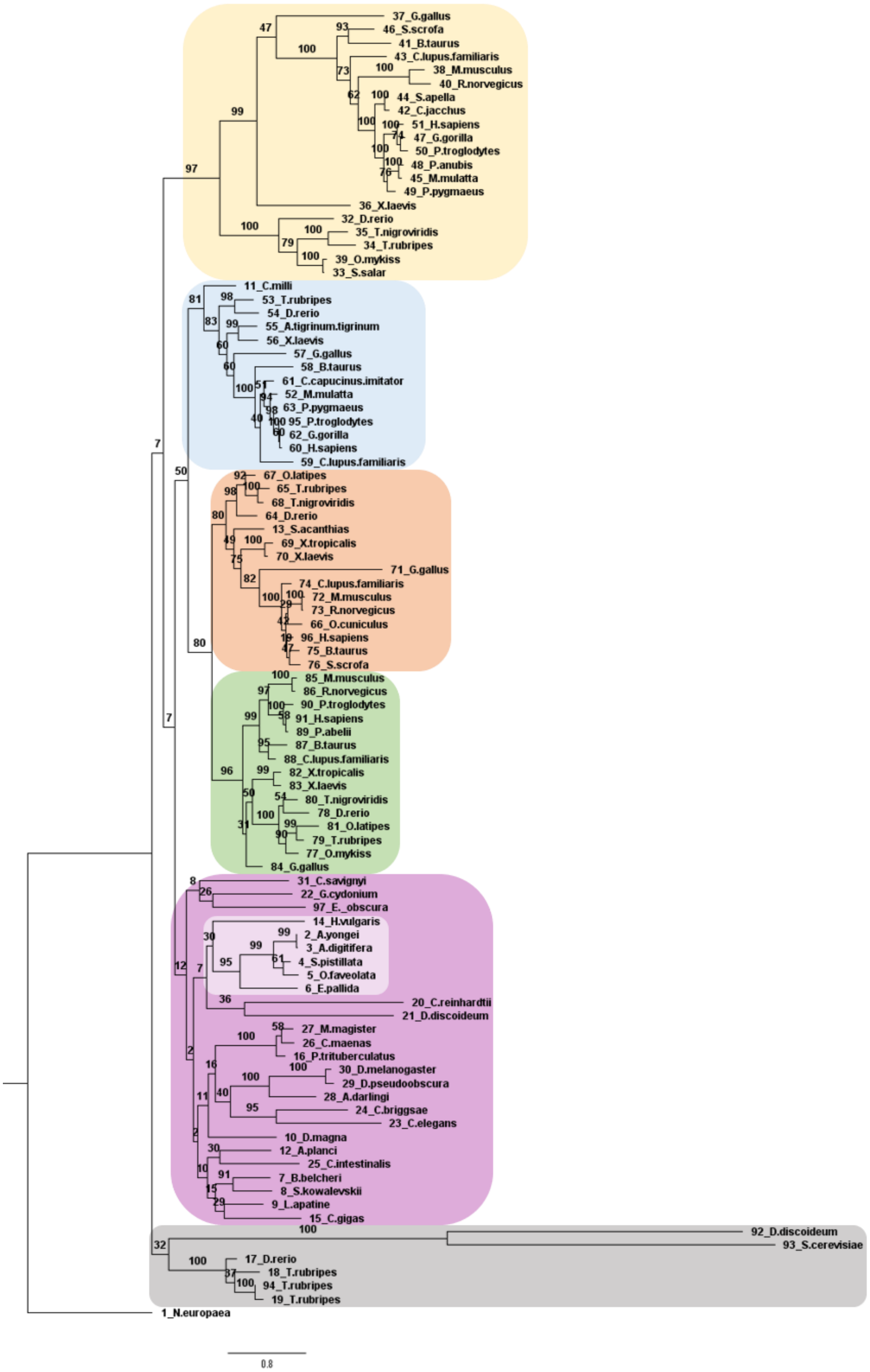

37_G.gallus
46_S.scrofa
41_B.taurus
43_C.lupus.familiaris
38_M.musculus
40_R.norvegicus
44_S.apella
42_C.jacchus
51_H.sapiens
47_G.gorilla
50_P.troglodytes
48_P.anubis
45_M.mulatta
49_P.pygmaeus
36_X.laevis
32_D.rerio
35_T.nigroviridis
34_T.rubripes
39_O.mykiss
33_S.salar
11_C.milli
53_T.rubripes
54_D.rerio
55_A.tigrinum.tigrinum
56_X.laevis
57_G.gallus
58_B.taurus
61_C.capucinus.imitator
52_M.mulatta
63_P.pygmaeus
95_P.troglodytes
62_G.gorilla
60_H.sapiens
59_C.lupus.familiaris
67_O.latipes
65_T.rubripes
68_T.nigroviridis
64_D.rerio
13_S.acanthias
69_X.tropicalis
70_X.laevis
71_G.gallus
74_C.lupus.familiaris
72_M.musculus
73_R.norvegicus
66_O.cuniculus
96_H.sapiens
75_B.taurus
76_S.scrofa
85_M.musculus
86_R.norvegicus
90_P.troglodytes
91_H.sapiens
89_P.abelii
87_B.taurus
88_C.lupus.familiaris
82_X.tropicalis
83_X.laevis
80_T.nigroviridis
78_D.rerio
81_O.latipes
79_T.rubripes
77_O.mykiss
84_G.gallus
31_C.savignyi
22_G.cydonium
97_E._obscura
14_H.vulgaris
2_A.yongei
3_A.digitifera
4_S.pistillata
5_O.faveolata
6_E.pallida
20_C.reinhardtii
21_D.discoideum
27_M.magister
26_C.maenas
16_P.trituberculatus
30_D.melanogaster
29_D.pseudoobscura
28_A.darlingi
24_C.briggsae
23_C.elegans
10_D.magna
12_A.planci
25_C.intestinalis
7_B.belcheri
8_S.kowalevskii
9_L.apatine
15_C.gigas
92_D.discoideum
93_S.cerevisiae
17_D.rerio
18_T.rubripes
94_T.rubripes
19_T.rubripes
1_N.europaea
0.8

**Supplementary Figure 1:** Maximum likelihood tree of the *Acropora yongei* Rh protein in relation to invertebrate and vertebrate Rh proteins. Rh family subgroups are denoted by color: Rh30 (yellow), Rhag (blue), Rhbg (orange), Rhcg (green), Rhp1 (light and dark purple), and Rhp2 (grey) as per (4). Cnidarian Rh proteins are highlighted within the Rhp1 subgroup (light purple). Amino acid sequences were aligned using MUSCLE and a maximum likelihood tree was generated using RAxML (500 bootstraps, PROTGAMMA model of rate heterogeneity, GTR substitution model). Sequences and the outgroup (*Nitrosoma europaea* Rh) are sourced from (4) with additional sequences identified by NCBI BLAST search. Sequence accession numbers are provided in Supplementary Table 1.

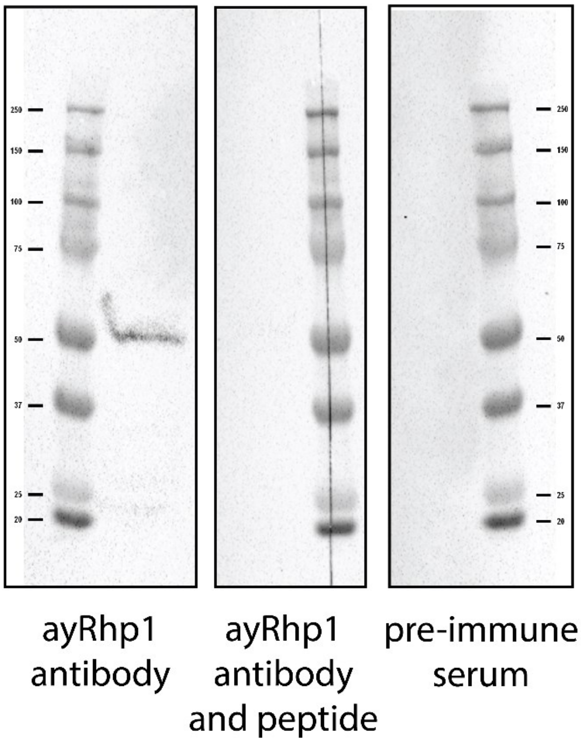


**Supplementary Figure 2:** Western Blot validation of the custom anti-ayRhp1 antibody. Membranes were labeled with anti-ayRhp1 primary antibody, primary antibody with excess peptide, or pre-immune serum at an equal concentration to primary antibody alone.

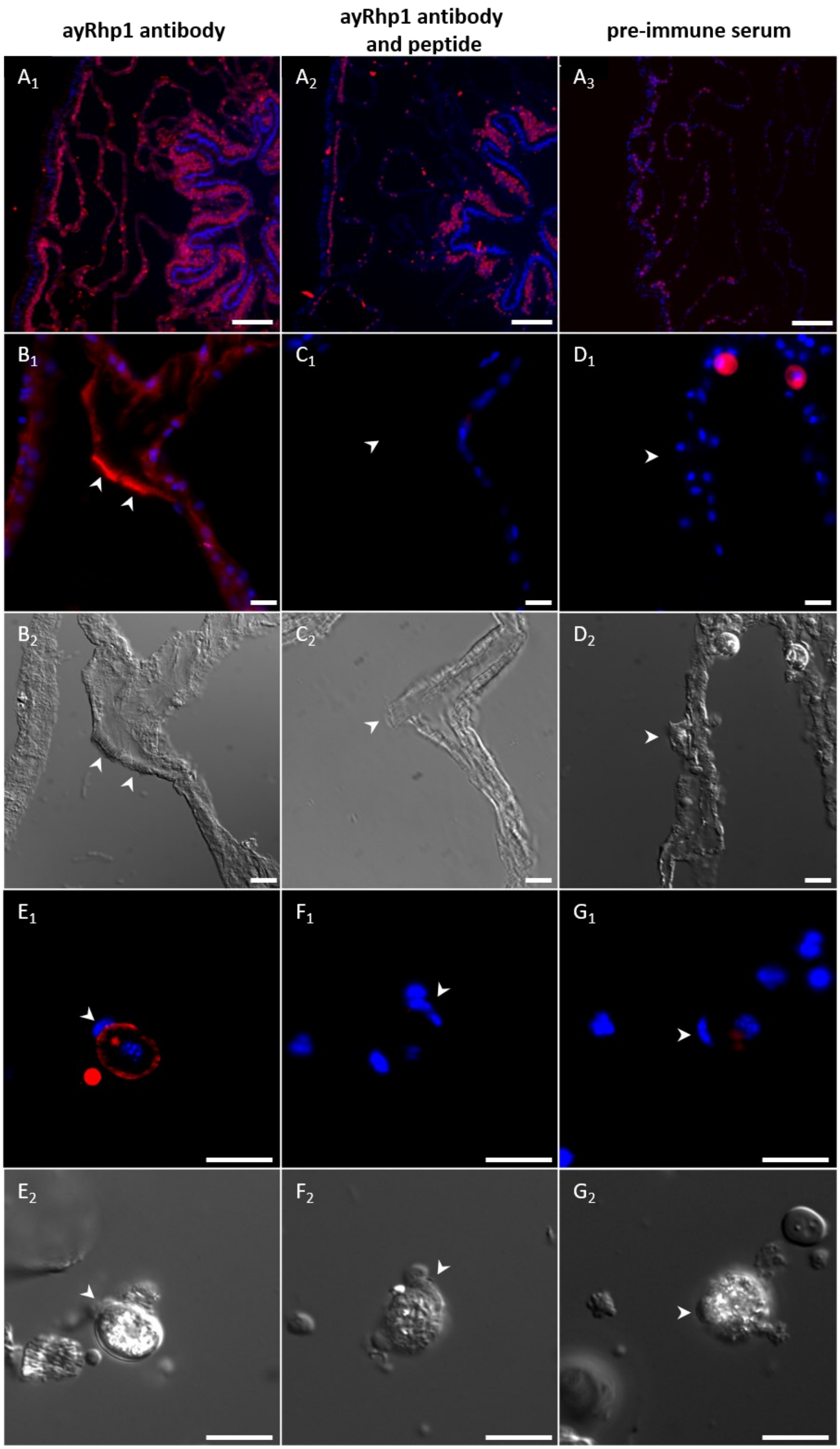
ayRhp1 antibody
ayRhp1 antibody and peptide
pre-immune serum
A1
A2
A3
B1
C1
D1
B2
C2
D2
E1
F1
G1
E2
F2
G2

**Supplementary Figure 3:** Immunohistochemistry validation of the custom anti-ayRhp1 antibody. Tissue sections and isolated cells were incubated with anti-ayRhp1 primary antibody, primary antibody with excess peptide, or pre-immune serum at an equal concentration to primary antibody alone. Anti-ayRhp1 antibody signal dissipates with peptide incubation and is absent in pre-immune serum incubations. **($A_{1-3}$)** Overview images of tissue sections. **($B_1$-$D_1$)** Desmocytes in tissue sections. **($E_1$-$G_1$)** Isolated coral host cells containing algal symbionts. **($B_2$-$G_2$)** corresponding brightfield differential interference contrast images. Nuclei are shown in blue, ayRhp1 in red. White arrowheads mark corresponding locations in epifluorescence and brightfield images. Scale bar = 100 μm **(A)** or 10 μm **(B-G)**.

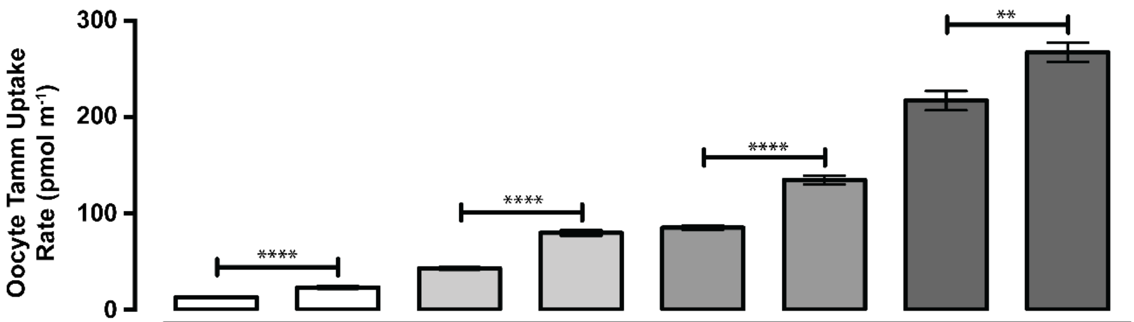


| | | | | | | | | |
|---|---|---|---|---|---|---|---|---|
| | SHAM | ayRhp1 | SHAM | ayRhp1 | SHAM | ayRhp1 | SHAM | ayRhp1 |
| pH | 6.5 | 6.5 | 7.5 | 7.5 | 8.5 | 8.5 | 7.5 | 7.5 |
| Tamm (µM) | 1,000 | 1,000 | 1,000 | 1,000 | 1,000 | 1,000 | 10,000 | 10,000 |
| $[NH_3]$ (µM) | 1.8 | 1.8 | 17.4 | 17.4 | 150.5 | 150.5 | 174.0 | 174.0 |
| $[NH_4^+]$ (µM) | 998.2 | 998.2 | 982.6 | 982.6 | 849.5 | 849.5 | 9826 | 9826 |

**Supplementary Figure 4:** Effect of $[NH_3]$ on total ammonium (Tamm) uptake rate in *Xenopus* oocytes expressing ayRhp1. Data shows mean ± S.E.M. of 6-14 oocytes; * denote significant differences (unpaired t-test; ***p<0.01; ****p<0001*).

**Supplementary Table 1:** NCBI accession numbers for sequences used in phylogenetic analysis (Fig. S1).

| ID | Accession Number | Organism |
|---|---|---|
| 1 | AY377923.1 | Nitrosomonas europaea |
| 2 | MH025799 | Acropora yongei |
| 3 | XP_015769291.1 | Acropora digitifera |
| 4 | XP_022795556.1 | Stylophora pistillata |
| 5 | XP_020600999.1 | Orbicella faveolata |
| 6 | KXJ18310.1 | Exaiptasia pallida |
| 7 | XP_019645061.1 | Branchiostoma belcheri |
| 8 | XP_006824549.1 | Saccoglossus kowalevskii |
| 9 | XP_013381459.1 | Lingula anatine |
| 10 | KZS19566.1 | Daphnia magna |
| 11 | AFK10779.1 | Callorhinchus milii |
| 12 | XP_022107421.1 | Acanthaster planci |
| 13 | AJF44128.1 | Squalus acanthias |
| 14 | XP_002167946.3 | Hydra vulgaris |
| 15 | EKC21768.1 | Crassostrea gigas |
| 16 | AHY27545.2 | Portunus trituberculatus |
| 17 | NP_571622.1 | Danio rerio |
| 18 | NP_001027816.1 | Takifugu rubripes |
| 19 | NP_001027817.1 | Takifugu rubripes |
| 20 | XP_001695464.1 | Chlamydomonas reinhardtii |
| 21 | XP_639042.1 | Dictyostelium discoideum |
| 22 | CAA73029.1 | Geodia cydonium |
| 23 | AAF97864.1 | Caenorhabditis elegans |
| 24 | XP_002636925.1 | Caenorhabditis briggsae |
| 25 | NP_001027959.1 | Ciona intestinalis |
| 26 | AAK50057.2 | Carcinus maenas |
| 27 | AEA41159.1 | Metacarcinus magister |
| 28 | ETN62951.1 | Anopheles darlingi |
| 29 | AAV40852.1 | Drosophila pseudoobscura |
| 30 | NP_001261434.1 | Drosophila melanogaster |
| 31 | AAY41910.1 | Ciona savignyi |
| 32 | NP_001019990.1 | Danio rerio |
| 33 | NP_001117044.1 | Salmo salar |
| 34 | NP_001027935.1 | Takifugu rubripes |
| 35 | AAY41905.1 | Tetraodon nigroviridis |
| 36 | NP_001084416.1 | Xenopus laevis |
| 37 | NP_989798.1 | Gallus gallus |
| 38 | AAC25123.1 | Mus musculus |
| 39 | AAP87367.1 | Oncorhynchus mykiss |

| | | |
|---|---|---|
| 40 | NP_071950.1 | Rattus norvegicus |
| 41 | NP_777137.1 | Bos taurus |
| 42 | AAF22442.1 | Callithrix jacchus |
| 43 | NP_001041501.1 | Canis lupus familiaris |
| 44 | AAF22501.1 | Sapajus apella |
| 45 | NP_001028136.3 | Macaca mulatta |
| 46 | NP_999543.1 | Sus scrofa |
| 47 | NP_001266526.1 | Gorilla gorilla |
| 48 | XP_003891396.1 | Papio anubis |
| ID | Accession Number | Organism |
| 49 | AAC94962.1 | Pongo pygmaeus |
| 50 | Q28813.2 | Pan troglodytes |
| 51 | P18577.2 | Homo sapiens |
| 52 | NP_001027987.1 | Macaca mulatta |
| 53 | NP_001032956.1 | Takifugu rubripes |
| 54 | NP_998010.1 | Danio rerio |
| 55 | AAV28818.1 | Ambystoma tigrinum tigrinum |
| 56 | XP_018121493.1 | Xenopus laevis |
| 57 | NP_989795.1 | Gallus gallus |
| 58 | NP_776596.1 | Bos taurus |
| 59 | NP_001104238.1 | Canis lupus familiaris |
| 60 | AHY04440.1 | Homo sapiens |
| 61 | XP_017354151.1 | Cebus capucinus imitator |
| 62 | NP_001266499.1 | Gorilla gorilla |
| 63 | AAG00305.1 | Pongo pygmaeus |
| 64 | NP_956365.2 | Danio rerio |
| 65 | NP_001027818.1 | Takifugu rubripes |
| 66 | NP_001075605.1 | Oryctolagus cuniculus |
| 67 | NP_001098561.1 | Oryzias latipes |
| 68 | Q3BBX8.1 | Tetraodon nigroviridis |
| 69 | NP_001011175.1 | Xenopus tropicalis |
| 70 | Q69D48.1 | Xenopus laevis |
| 71 | AAN34362.1 | Gallus gallus |
| 72 | AAF19371.1 | Mus musculus |
| 73 | NP_898877.1 | Rattus norvegicus |
| 74 | NP_001003017.2 | Canis lupus familiaris |
| 75 | NP_777148.1 | Bos taurus |
| 76 | NP_999161.1 | Sus scrofa |
| 77 | NP_001117995.1 | Oncorhynchus mykiss |
| 78 | AAM90586.1 | Danio rerio |
| 79 | NP_001027934.1 | Takifugu rubripes |
| 80 | Q3BBX7.1 | Tetraodon nigroviridis |
| 81 | NP_001116374.1 | Oryzias latipes |

| | | |
|---|---|---|
| 82 | XP_012814245.1 | Xenopus tropicalis |
| 83 | Q5U4V1.1 | Xenopus laevis |
| 84 | NP_001004370.1 | Gallus gallus |
| 85 | AAF19373.1 | Mus musculus |
| 86 | NP_898876.1 | Rattus norvegicus |
| 87 | NP_776597.1 | Bos taurus |
| 88 | NP_001041487.1 | Canis lupus familiaris |
| 89 | XP_002825848.1 | Pongo abelii |
| 90 | XP_016782540.1 | Pan troglodytes |
| 91 | NP_057405.1 | Homo sapiens |
| 92 | BAB39709.1 | Dictyostelium discoideum |
| 93 | X77608.1 | Saccharomyces cerevisiae |
| 94 | AAU81656.1 | Takifugu rubripes |
| 95 | NP_001009033.1 | Pan troglodytes |
| 96 | AF193807.1 | Homo sapiens |
| 97 | AJO26542.1 | Erpobdella obscura |
